# NanoCT-confocal mapping of mouse ovaries reveals lifespan-persistent symmetric organizational rules of primary-to-preovulatory follicles

**DOI:** 10.1101/2025.10.29.685326

**Authors:** Giulia Fiorentino, Andrea Fantinato, Annapaola Parrilli, Tommaso Marzi, Cesare Alippi, Riccardo Bellazzi, Laura Rienzi, Filippo Maria Ubaldi, Alberto Vaiarelli, Valeria Merico, Paola Rebuzzini, Danilo Cimadomo, Silvia Garagna, Maurizio Zuccotti

## Abstract

Mammalian ovaries are organized into follicles integrated with the surrounding tissue, collectively shaping an architecture continuously remodeled through cycles of growth and elimination. Combining nano-Computed Tomography with 3D digital modeling, we deciphered spatial rules governing follicle organization in the mouse ovary across lifespan. We find that primary to preovulatory follicles adopt a quantitatively symmetric arrangement along the anterior-posterior and dorsal-ventral axes, established during prepuberty before secondary follicle vascularization and preserved through adulthood and aging, regardless of follicle health or atresia. A novel nanoCT-confocal pipeline revealed that follicles enclosing oocytes with transcriptionally-active or transcriptionally-inactive chromatin states follow divergent fates of growth, atresia, or developmental competence, yet retaining a symmetric distribution. Mathematical modeling of follicle growth and elimination from prepuberty to adulthood identified transition rates between follicle stages that support gonadal symmetry. These findings suggest an evolutionarily optimized developmental program ensuring follicle recruitment and high ovulatory output, a hallmark of polytocous species.

## INTRODUCTION

The activity of many organs relies on specialized functional units that integrate with the surrounding tissue, interact with one another, and occupy a defined spatial location, contributing to shape a specific functional architecture. This organization, together with distinct molecular and physiological profiles, determines how an organ maintains homeostasis or transitions to dysfunction. To understand how the organ’s three-dimensional (3D) organization coordinates its activities requires knowing not only its molecular context, but also the accurate, quantitative, and spatial map of its functional units and to be able to follow their changes over time.

The mammalian ovary is an example of this condition, perhaps the most remarkable, as it undergoes continuous restructuring of its internal architecture during regular activity. Follicles, the functional units of the female gonad, undertake differentiation that profoundly transforms them and reshapes the ovary’s organization. During folliculogenesis, which is coordinated by endocrine and paracrine signaling pathways and biomechanical cues, follicles change in both size and function, coordinating growth and acquisition of oocytes developmental competence [1].

In the mouse, a polytocous species that ovulates several oocytes per reproductive cycle, folliculogenesis begins with primordial follicles, in which a small oocyte is either naked (Type 1, T1) or surrounded by a single layer of flattened pregranulosa cells (T2). They then advance to primary follicles (T3), marked by a cuboidal granulosa layer and the onset of zona pellucida (ZP) deposition; the secondary follicles with two (T4) or multiple (T5) granulosa layers and theca-cell recruitment; the early-antral (T6) and antral (T7) follicles; and finally, the preovulatory (T8) [2]. Across the T1-T8 progression, atresia eliminates non-viable follicles and leaving zona pellucida remnants (ZPRs), persistent glycoprotein shells that mark completed degeneration [3].

Although the progression of folliculogenesis, from primordial activation to preovulatory maturation is well described, its spatiotemporal organization inside the ovary remains poorly characterized. Key questions about how follicles are spatially arranged, how their distribution changes over time, and how growth and atresia evolve in the 3D context and across life remain unsolved.

Efforts to address these questions have employed a range of methods to reconstruct the ovary’s 3D organization [4,5]. Traditional histology, while widely used, is prone to misalignment and loss of spatial context [6]. Confocal and light-sheet fluorescence microscopy, though allowing the identification of molecular aspects, suffer from limited resolution in the depth axis and require extensive tissue clearing [7,8]. Micro Computed Tomography (microCT), which we used in a pilot study on adult mouse ovaries [9], preserves the native tissue architecture but provides only moderate resolution, limiting its capacity to resolve fine morphological details.

Here, to uncover the spatial logic of folliculogenesis, we leveraged nanoCT’s high-resolution isotropic imaging to develop a novel pipeline capable of integrating structural, spatial, viability (healthy vs. atretic), and transcriptional dimensions. With a voxel size reaching 250 nm and an optimized fixation, contrast-enhancement and embedding protocol, we mapped follicles from the primary (T3) to the preovulatory (T8) stage. Ovaries collected across sixteen developmental timepoints, from infancy (1-10 days post birth, dpb), prepuberty (11-25 dpb), adulthood (2-4 months post birth, mpb) and advanced age (9-18 mpb), yielded over fifty volumetric datasets capturing the ovary’s dynamic maturation landscape. Each sample was digitally reconstructed, annotated, and bisected along the anterior-posterior (A-P) or dorsal-ventral (D-V) central planes, enabling both cross-sectional division and axis-wise volumetric analysis to map follicles, quantify their number and viability, and establish a mathematical framework for estimating rules underlying their spatial distribution. We then devised a nanoCT-confocal imaging strategy that mapped T3-T8 follicles by oocyte’s chromatin configuration, distinguishing transcriptionally active from silent states and developmental competency.

The results of this study offer the first multimodal 3D atlas of the mouse ovary that captures temporal dynamics and a striking, spatially ordered, symmetry of follicle populations, set early in life, maintained across the reproductive lifespan, and largely independent of follicle healthy or atretic status.

## RESULTS

### High-resolution nanoCT and 3D modeling reveal morphological features of folliculogenesis and atresia

To enable comprehensive *in situ* mapping of ovarian follicles, we established a three-step pipeline integrating nanoCT imaging, expert-based annotation, and interactive 3D digital modeling. After initial setting on 25 dpb ovaries, this approach was applied across the female reproductive lifespan.

To achieve high-resolution 3D visualization and classification of ovarian follicles, we first established a contrast-enhanced nanoCT imaging protocol optimized for deep tissue penetration and differential radiopacity, with an isotropic voxel size of 250 nm (Video S1; Figure 1A). NanoCT classification of primary (T3) to preovulatory (T8) follicles was based on morphometric parameters including follicular diameter, granulosa cell layering, presence of a ZP space, and oocyte features, as detailed in Figure 1B, C. Beyond staging healthy follicles, nanoCT also enabled identification of seven distinct morphological hallmarks of follicular atresia, consistent with those described in earlier histological studies [3] and detailed in Figure 1D. Thus, the nanoCT-reconstructed mouse ovary provides a comprehensive 3D framework for precise staging of folliculogenesis and detailed characterization of atresia.

**Figure 1.**
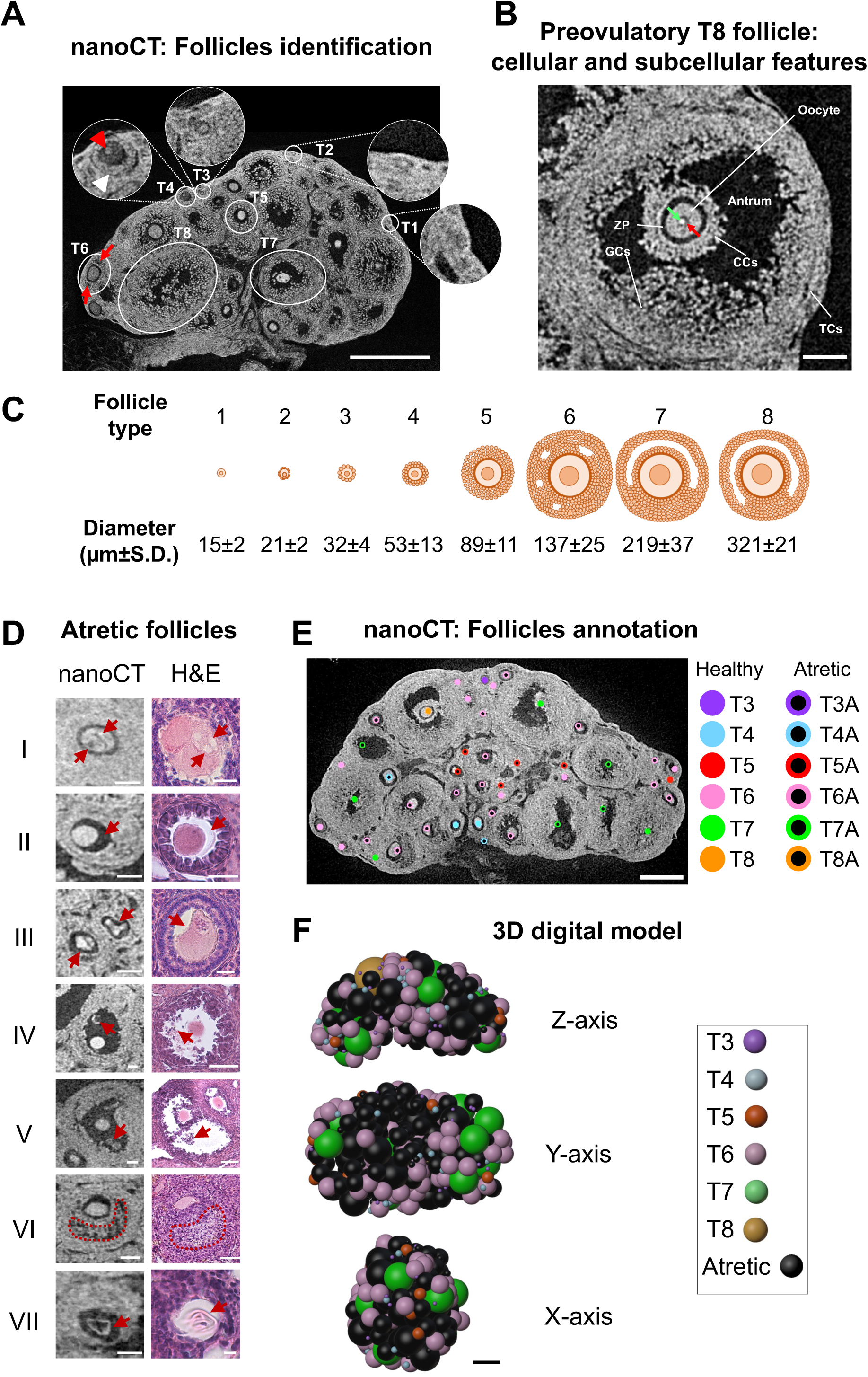
Contrast-enhanced nanoCT enables 3D follicle staging, atresia characterization, and digital reconstruction of ovarian architecture (A) Representative nanoCT image of the equatorial section of a 25 dpb ovary showing all the different T1-T8 follicle types. Despite high-resolution imaging, the quantification of primordial T1 and T2 follicles in the cortical periphery was hindered by their abundance, estimated at ∼5,000 in young adults, and a few hundred in 12 mpb mice [27–29]. T3 follicles display a single layer of cuboidal granulosa cells encasing a small oocyte (inset). In T4 follicles, nanoCT delineate a double granulosa layer (white arrowhead), the ZP, and a nucleated oocyte (red arrowhead). Preantral T5 follicles exhibit multilayered granulosa cells without antrum formation, whereas early antral T6 follicles show incipient antral cavities (red arrows) bordered by defined granulosa and theca layers. T7 follicles show a prominent antrum with a discernible cumulus oophorus. Insets show enlargements of T1 to T4 follicles. Bar, 200 µm. (B) nanoCT image enlargement of a fully grown T8 follicle displaying distinct theca (TCs) and granulosa (GCs) cell layers, antrum cavity, cumulus cells (CCs), ZP, and an oocyte with visible nucleus (red arrow) and nucleolus (green arrow). Bar, 100 µm. (C) Follicle classification (T3-T8) based on morphometric features including follicular diameter, granulosa cell layering, ZP presence, and oocyte morphology [2,9]. (D) Temporal sequence of seven morphological hallmarks of follicular atresia identified by nanoCT: (I) oocytes exhibiting small cytoplasmic cavities (red arrows); (II) granulosa cell retraction from the ZP (red arrows); (III) structurally disorganized degenerating oocytes (red arrows); (IV) degenerating oocytes with sparse pyknotic cumulus cells (red arrows); (V) apoptotic granulosa aggregates within the antral cavity (red arrows); (VI) presence of fibroblast-like cells into the antrum (red dots contour); (VII) complete oocyte loss with a retained ZPR (red arrows). Bar nanoCT images: I-VII, 50 µm; Bar histology images: I-III and VII, 20 µm; IV-VI, 50 µm. (E) Follicle identification and annotation is performed on the full nanoCT dataset using Fiji, enabling 3D mapping of healthy (T3-T8) and atretic (T3A-T8A) follicles. Bar, 250 µm. (F) Digital 3D ovarian model reconstructed using Blender with follicle position, size, and status (healthy vs. atretic) displayed. These interactive models allow rotation along x, y, and z axes and toggling on/off of follicles by maturational stage or health/atretic status. Bar, 200 µm.

Then, to facilitate accurate 3D visualization and spatial analysis of the ovary across the full nanoCT image stack, we processed the datasets using the open-source software Fiji [10]. This allowed precise and robust identification, annotation and counting of all healthy (T3-T8) and atretic (T3A-T8A) follicles (Figure 1E; Video S2).

Next, to enable the exploration of the ovarian architecture we used the open-source platform Blender to produce 3D digital models. A custom Python script mapped the x, y, and z coordinates of follicles center, and another script imported csv files containing the coordinates, average follicle sizes (Figure 1C), and assigned color codes, with atretic follicles displayed in black. Once reconstructed, each ovary model could be interactively rotated along the x, y, or z axis (Figure 1F; Video S3A). Single or grouped follicles, classified by developmental stage or by healthy/atretic status, could be independently toggled on or off, offering a user-friendly, customizable interface for both qualitative and quantitative analyses (Video S3B).

To validate the fidelity of the 3D digital model, we compared follicle counts along the anatomical A-P and D-V axes obtained using expert-based and Blender-based approaches. A third custom Python script automatically sectioned the ovary (Figure S1A) and categorized follicles along the A-P and D-V axes. A side-by-side comparison showed high concordance in follicle number and spatial distribution (Figure S1B), confirming the accuracy of the automated model despite its simplified geometric assumptions, which treated follicles as spheres of uniform diameter. This approach enabled the generation of simplified digital infant, prepubertal and adult ovaries, providing an interactive means to investigate follicular dynamics in situ and intuitively explore emergent spatial arrangements across ages in silico.

### During infancy and prepuberty the ovary builds an ordered spatial distribution of both healthy and atretic follicles

To investigate when growing T3-T8 follicles first emerge, and to quantitatively map their distribution during infancy and prepuberty (Figure 2A) we used the nanoCT-based classification framework described above.

**Figure 2.**
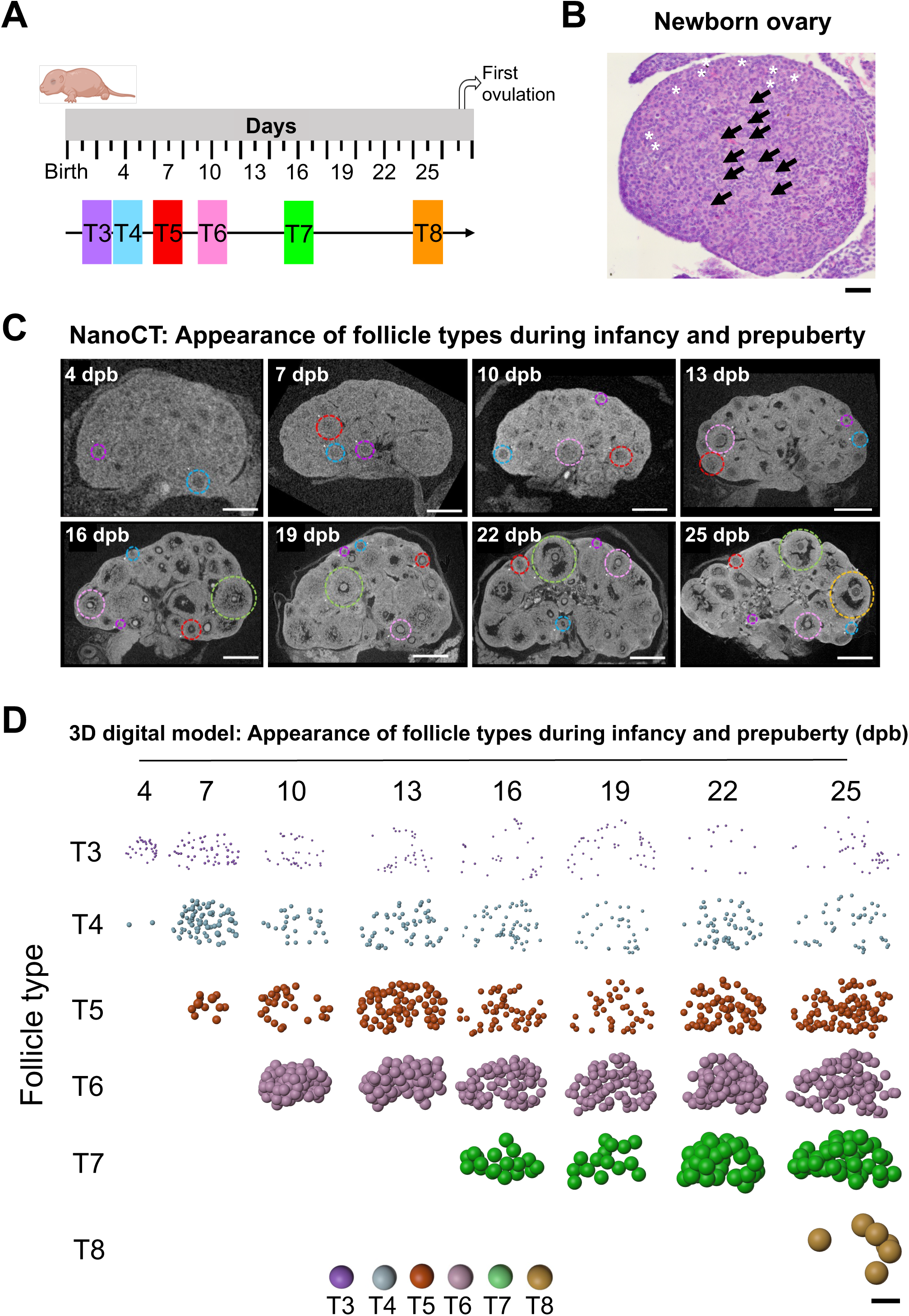
Spatiotemporal mapping of follicle growth in the infant and prepubertal ovary (A) Overview of the developmental stages analyzed by nanoCT (top) and timeline of T3-T8 follicle first appearance across infancy (1-10 dpb) and prepuberty (11-25 dpb) (bottom). (B) Representative histological equatorial section of a newborn ovary showing abundant T1-T2 primordial follicles throughout the gonad (white asterisks) and centrally located clusters of T2-T3 follicles (black arrows), indicating a centrally initiated first wave of folliculogenesis. Bar, 50 µm. (C) NanoCT-based classification and mapping of T3-T8 follicles across infant and prepubertal stages. The nanoCT images show the temporal emergence of T3 with T4, T5, T6, T7 or T8 follicles captured at 4-, 7-, 10-, 16-, and 25 dpb, respectively. Bar, 4-10 dpb, 100 µm; 13-19 dpb, 200 µ; 22 dpb, 250 µm; 25 dpb, 300 µm. (D) 3D digital model illustrating the progressive spatial distribution and maturation of T3-T8 follicles, with fully grown T8 follicles detected by 25 dpb. Bar, 350 µm.

At birth, the mouse ovary is a spherical structure ∼0.5 mm in diameter that matures into an ovoid, bean-shaped organ (∼1×2×1 mm) by 25 dpb. Due to limited imaging contrast in newborn ovaries, individual follicles could not be resolved by nanoCT; therefore, classical histological analysis was employed for this developmental stage only. Thousands of T1-T2 primordial follicles were distributed throughout the gonad, with a centrally located cluster of T2-T3 follicles (Figure 2B). This spatial arrangement confirms that the first wave of folliculogenesis initiates centrally [11,12], when T3 follicles acquire a cuboidal granulosa cell layer. NanoCT imaging detected T4 follicles by 4 dpb (Figure 2C), with T5 and T6 emerging at 7 and 10 dpb, respectively. By 16 dpb, T7 follicles were present, and by 25 dpb, fully grown preovulatory T8 had appeared. Figure 2D shows a 3D digital model of the appearance of T3-T8 follicles.

To map the onset of atresia, we quantified healthy and atretic follicles across infant and prepubertal development. Total follicle number increased gradually ∼3-fold from 4 to 25 dpb (Figure 3A). Morphological atretic markers were absent between 4 and 16 dpb but appeared thereafter, stabilizing at ∼31-49% of all follicles between 19 and 25 dpb (Figure 3B, C).

**Figure 3.**
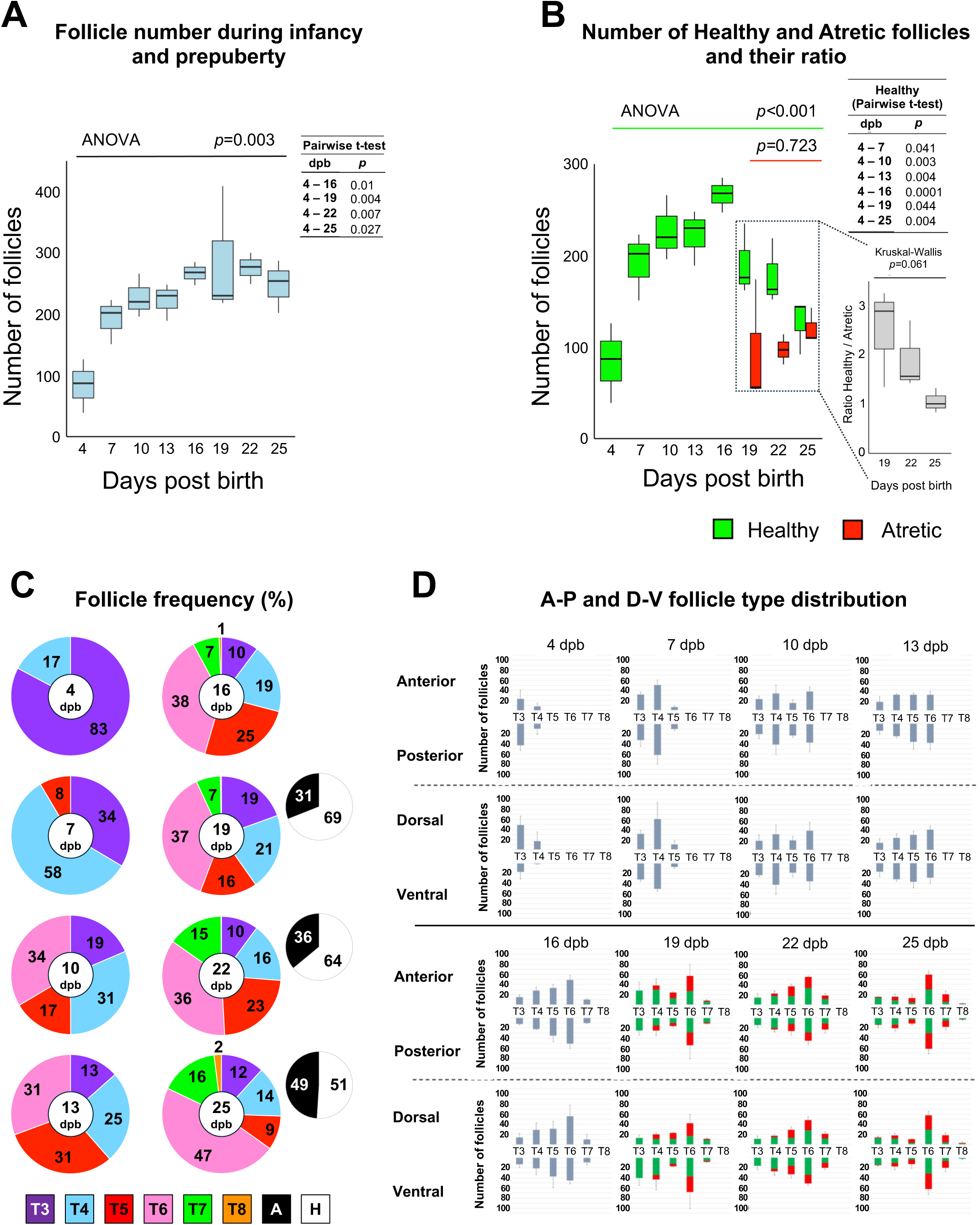
Number of follicles and onset and spatial distribution of follicular atresia in the infant and prepubertal ovary (A) Box plot showing the total number of follicles quantified by nanoCT across infant and prepubertal development (4-25 dpb). Statistical analysis describes a highly significative∼3-fold increase in total follicle count. (B) Box plot showing that atresia emerges from 19 dpb and stabilizes at ∼31-49% of total follicles, with no statistically significant difference between 19, 22 and 25 dpb (right inset). (C) Follicle type frequency across infant and prepubertal ages, highlighting T6 follicles as the most abundant type. (D) Spatial distribution of T3-T8 healthy and atretic follicles across A-P and D-V regions during infancy and prepuberty, showing symmetric allocation without regional enrichment or depletion.

Next, we asked which follicle type contributed most to the observed increase, and which was most affected by atresia. Healthy and atretic follicle counts were singled out into single follicle types (Figure S2A). The atretic pool was largely composed of T5 and T6 follicles, with T6 accounting for nearly half of all atretic events (Figure S2B), reflecting its numerosity within the growing pool (Figure 3C). In contrast, T3 follicles were rarely affected, appearing occasionally atretic at 25 dpb (Figure S2A).

Then, to assess spatial organization, we oriented and virtually bisected the nanoCT datasets of whole ovaries along the A-P and D-V axes (Video S1), quantifying the distribution of each follicle type. Across all developmental stages, both healthy and atretic follicles were symmetrically distributed, with comparable numbers and types between the opposing halves of each axis, and no evidence of regional enrichment or depletion (Figure 3D). To investigate whether follicle health or atresia is associated with spatial localization over time, we generated in Blender a centered ellipsoid reference representing the inner 25% of ovarian volume, automatically coloring follicles grey when located within this region (Video S4). During ovarian development from 19 to 25 dpb, atretic follicles clustered toward the center, and by 25 dpb were significantly more centrally localized than their healthy counterparts (Figure S3A, B). Overall, these observations highlight a numerical preponderance of early antral T6 follicles and reveal an ordered spatial distribution of both healthy and atretic follicles, suggesting a centripetal shift in the 3D distribution of degenerating follicles during the late prepubertal window.

### Follicles enclosing oocytes with distinct transcriptional activity and healthy or atretic status display a symmetric ovarian organization

To further enrich the 3D reconstruction with functional insights, beyond identifying atretic follicles, we employed a co-registration approach that combined nanoCT mapping with DAPI-based confocal imaging. The latter was performed on serial 20 µm-thick histological sections from the same 25 dpb ovary, when the gonad has fully established its organization (Figure S4).

This novel multimodal method enabled classification of each T3-T8 follicle (Video S5A) based on healthy or atretic status (Figure 1D) and NSN or SN oocyte’s chromatin organization (Figure 4A), a marker of active or inactive transcriptional activity, respectively [13,14]. The 25 dpb ovaries contained a mean±S.D. of 247.7±42.9 follicles. Of these, only a small minority (1.3±1.4% *per* ovary) of healthy follicles lost their oocyte nucleus during histological processing and sectioning. An additional 21.8±5.5% could not be classified as NSN or SN because of advanced atresia and chromatin disorganization. Of these, the most abundant (∼50%) were atretic T6 follicles. As a result, 191±38.2 follicles, healthy and atretic, were classified as either NSN or SN (Figure 4B; Video S5B); approximately two-thirds displayed an NSN chromatin configuration (78.3±6.0% healthy; 21.7±6.0% atretic), while the remaining third exhibited an SN conformation (21.7±6.1% healthy; 78.3±6.1% atretic).

**Figure 4.**
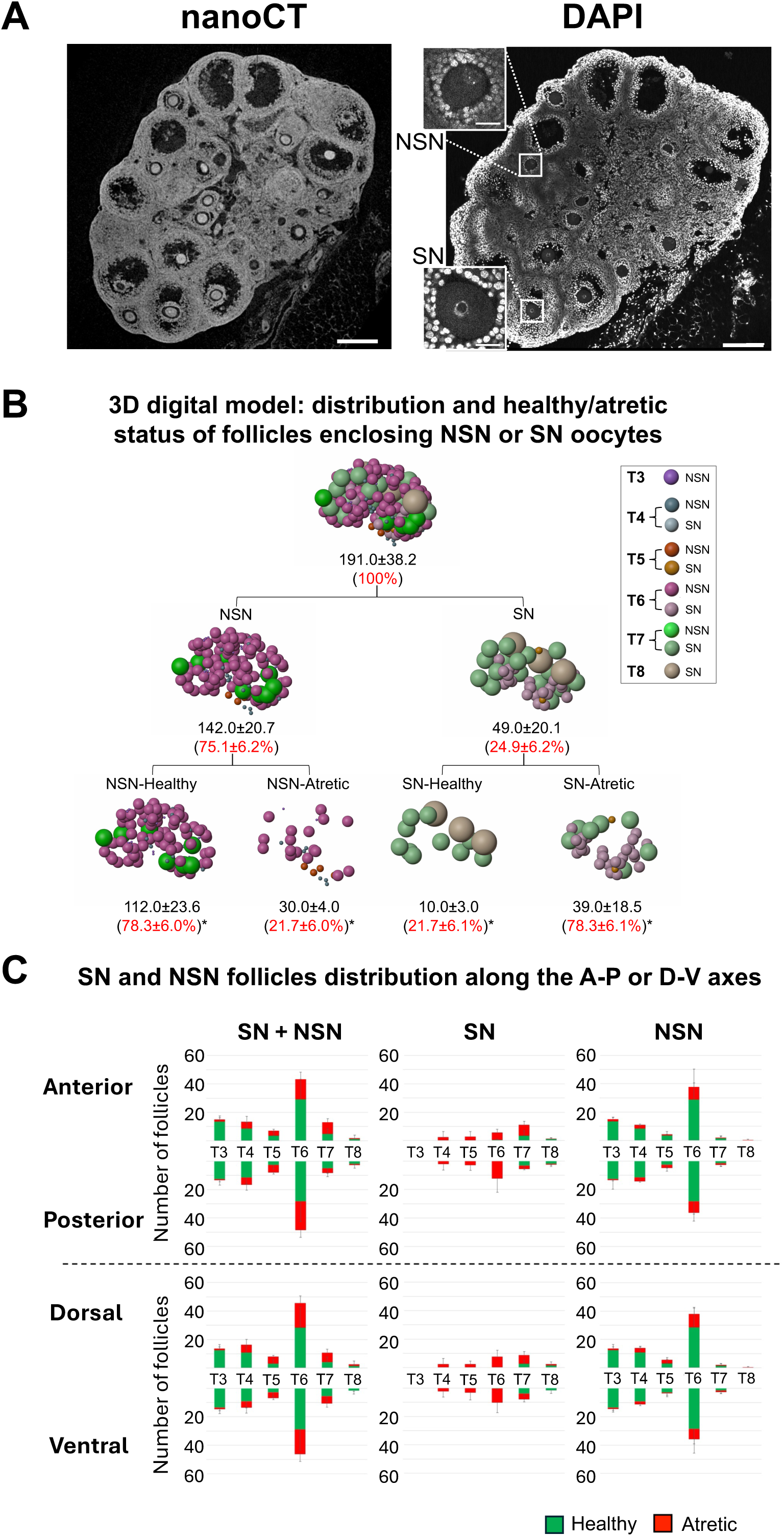
A nanoCT-confocal imaging pipeline enables functional mapping of follicles by healthy/atretic status and oocyte’s chromatin configuration (A) nanoCT-based 3D mapping (left) with DAPI-stained confocal imaging of serial 20 µm-thick sections (right) from the same 25 dpb ovary. T3-T8 follicles are classified by health status (nanoCT: healthy vs. atretic) and oocyte’s chromatin configuration (DAPI: NSN vs. SN). Bar: 200 µm. (B) Representative 3D digital model of a 25 dpb ovary allowing quantification (Mean±S.D. and %±S.D.), spatial mapping and visual observation of T3-T8 follicles depending on their healthy/atretic status and NSN/SN chromatin configuration. *: % relative to the preceding category. (C) Spatial distribution of T3-T8 follicles along the A-P or D-V axes, based on their healthy/atretic status and NSN or SN oocyte’s chromatin configuration. The histograms show an even distribution of all follicle types, healthy or atretic, SN or NSN, across planes.

To assess the spatial distribution of SN and NSN follicles within the ovary, the nanoCT datasets were virtually bisected along the A-P and D-V axes, revealing an even distribution of all follicle types, healthy or atretic, NSN or SN, across both planes (Figure 4C). In summary, NSN oocytes were abundantly present from the primary T3 to the antral T6, and in small numbers in the subsequent T7 and T8, the latter associated only to atresia. Instead, SN oocytes first appeared in preantral T4 through T6 follicles, where they were consistently associated with morphological signs of atresia, while in T8 they were found enclosed in healthy follicles. Notably, when considering only healthy follicles, NSN oocytes were present from T3 to T7, whereas SN oocytes were restricted to T7 and T8.

### Spatiotemporal dynamics of folliculogenesis during adulthood and aging confirm persistent ovarian symmetry despite a decline in follicle number

To investigate how follicle abundance, health status, and spatial organization evolve across reproductive aging, we quantitatively assessed healthy and atretic follicle populations from early adulthood to advanced age (Video S6).

NanoCT equatorial sections of 2-18 mpb ovaries revealed distinct age-associated changes in ovarian architecture (Figure 5A). By 18 mpb extensive cystic structures [15] dominated the ovarian parenchyma (Figure 5A), indicating advanced tissue remodeling and limiting further structural resolution. Following puberty, the presence of CL confirmed completed folliculogenesis and active ovulatory cycles. On average, 23.9±15.6 CL *per* ovary were observed, exhibiting variable radiodensity, potentially reflecting different stages of luteal regression. Although we did not investigate this aspect further, as the present study focused on the follicular component, it clearly warrants future examination.

**Figure 5.**
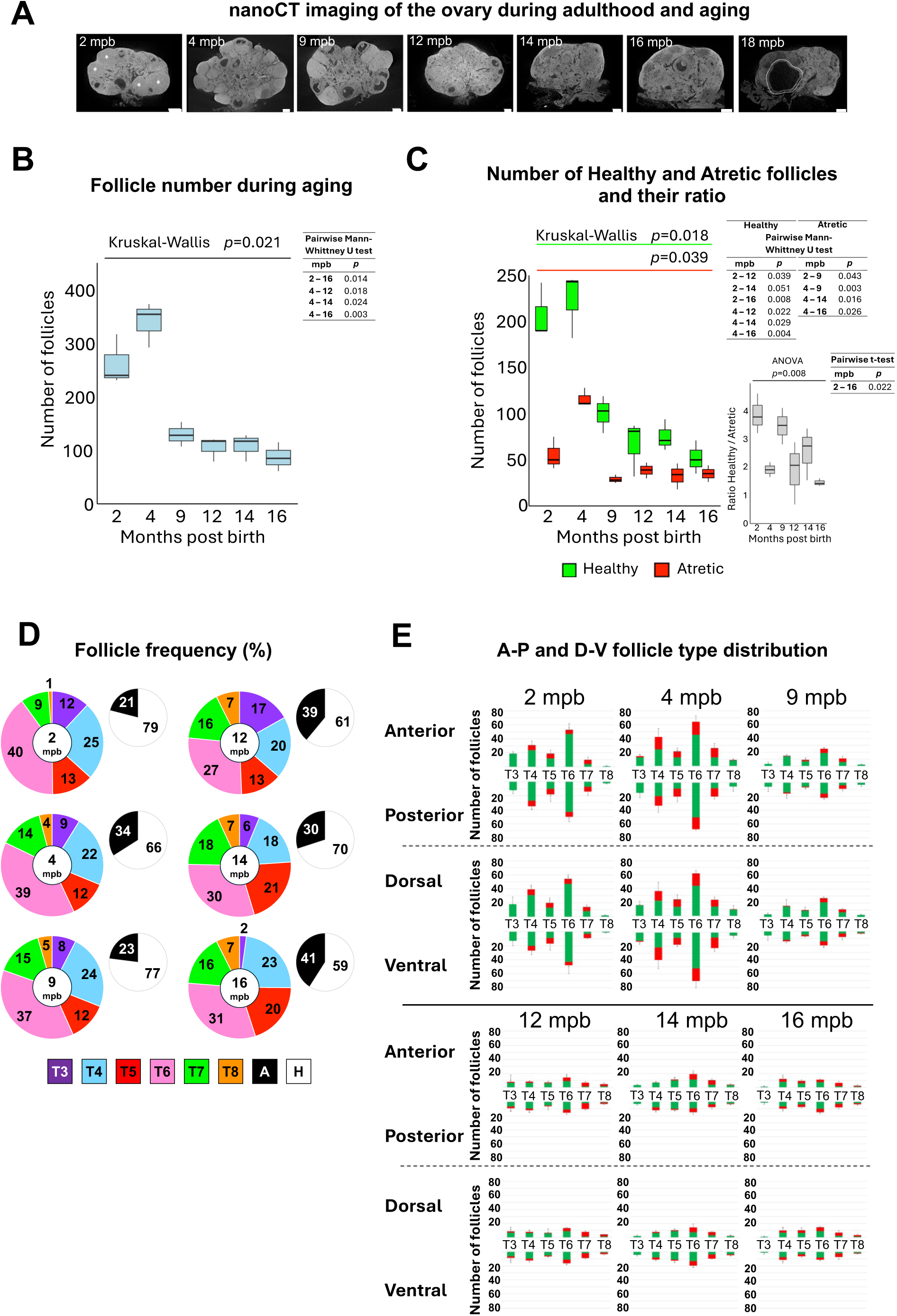
Age-associated changes in ovarian architecture, follicle dynamics, and spatial organization (A) Representative nanoCT equatorial sections of 2-18 mpb ovaries showing age-related structural changes. Adult ovaries contain CL (white asterisks), consistent with active ovulatory cycles, while 18 mpb ovaries exhibit extensive remodeling due to cysts formation. Bars: 250 µm. (B) Box plot of the total follicle number during adulthood and aging, showing a sharp 50% decline from the adult (2-4 mpb) to the aging (9-16 mpb) ovary. (C) Box plots of the number of healthy and atretic follicles (left) and their ratio (right), showing that the proportion of healthy versus atretic follicles remained unchanged until 14 mpb and then declined significantly at the following 16 mpb age. (D) Frequency of follicle types across adulthood and aging. As during prepuberty, T6 is the most abundant follicle type. During aging atresia increases exceeding 40% by 16 mpb. (E) Spatial distribution of healthy and atretic follicles along the A-P and D-V axes across adulthood and aging, showing persistent symmetry despite the age-related decline in follicle abundance.

Total follicle counts remained stable from the latest prepubertal stages (19-25 dpb) through early adulthood (2-4 mpb) (Figure S2C), while the frequency of healthy follicles increased significantly (Figure S2D), mainly due to an increase of the number of T6 follicles. From 9 mpb, however, follicle numbers declined sharply by ∼50% and subsequently stabilized from 12 mpb into advanced age (Figure 5B). Whilst the ratio between healthy and atretic follicles remained unchanged until 14 mpb (Figure 5C), atretic T7 and T8 increased (Figure S2B), underlying a deterioration of the folliculogenetic process that becomes statistically evident by 16 mpb (Figure 5C), when the frequency of atretic follicles exceeds 40% (Figure 5D). Importantly, follicle type analysis (Figure S2A) identified a constant number of T3-T7 follicles across two time-windows: from late prepuberty through adulthood (22 dpb to 4 mpb), and, after drastic reduction in total follicle number, during the aging period (9-16 mpb).

Next, to determine whether ovarian aging affects follicles spatial distribution, we mapped the localization of healthy and atretic follicles within adult and aging ovaries. NanoCT analysis disclosed a uniform distribution along the A-P and D-V axes of healthy and atretic follicles between the two halves at any time point examined (Figure 5E). This spatial symmetry persisted despite the significative age-related decline in total follicle numbers. Together, these findings revealed that the spatial symmetry established during prepuberty is preserved in adult ovaries at 2 and 4 mpb, corresponding to peak fertility. Although follicle abundance declines markedly with age, their anatomical distribution remains largely isotropic, indicating that ovarian aging does not alter the spatial organization of the follicular niche.

To track the fate and spatial dynamics of glycoprotein-rich ZPRs, we analyzed PAS-hematoxylin-stained histological sections of the same nanoCT samples confirming the presence of ZPRs in corresponding regions (Figure S5A). ZPRs were absent in prepubertal ovaries, first appeared by 2 mpb, at ∼200/ovary, and showed a highly significant 10-fold increase by 12 mpb (Figure S5B), predominantly accumulating within the 25% inner ellipsoid (Figure S5C), before declining in later ages (Figure S5B). Together, these findings demonstrate that ZPRs, whilst also scattered in the cortex, are progressively enriched towards the ovarian medulla (Video S6), suggesting regionally biased retention of follicular remnants during reproductive aging. To enable detailed visualization of age-related changes in follicle spatial organization, we generated 3D digital models of the adult ovary across 2-16 mpb (Figure 6).

**Figure 6.**
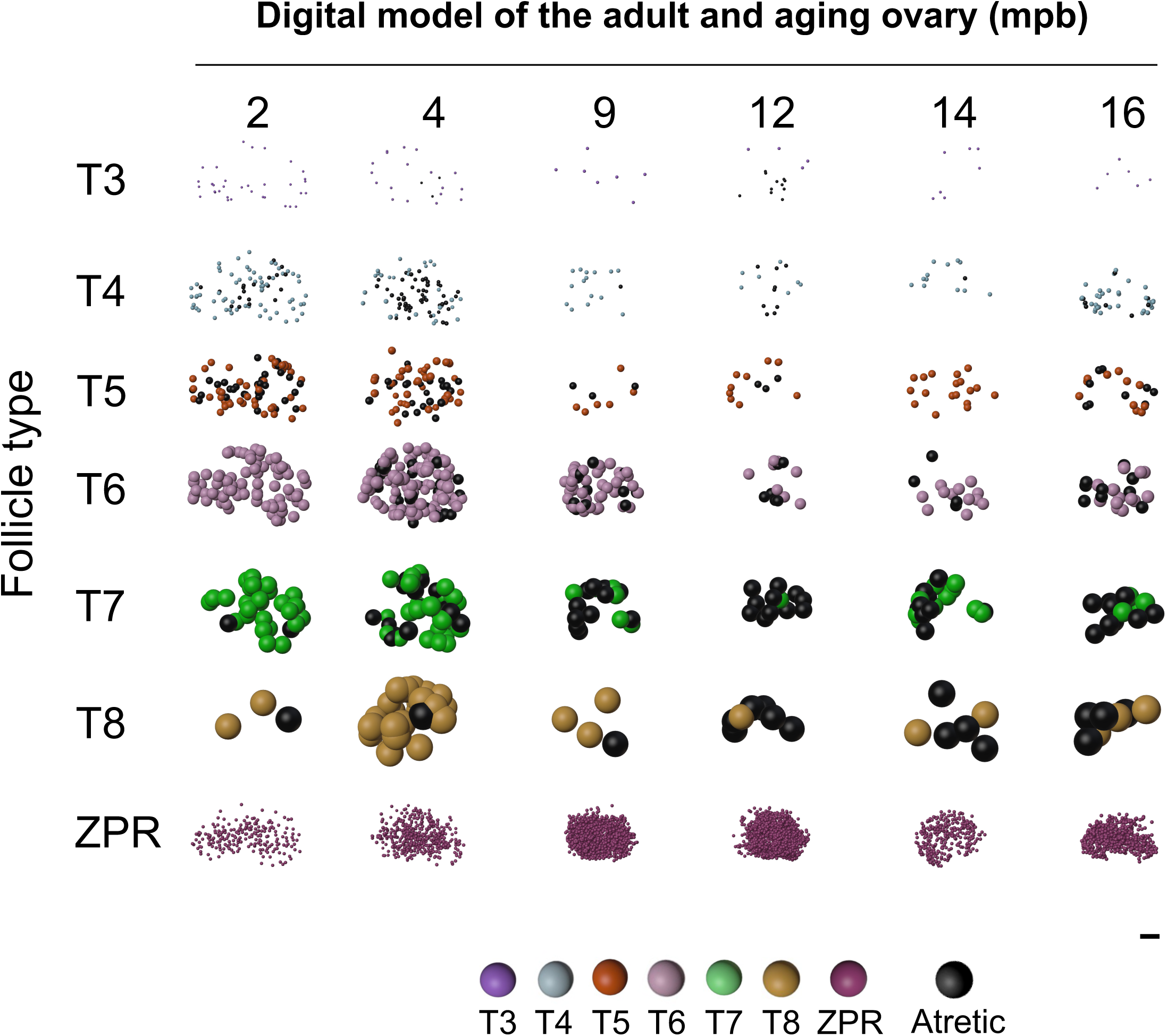
3D digital model illustrating the spatial distribution of T3-T8 follicles in adulthood and during aging A representative 3D digital reconstruction of 2-16 mpb ovaries generated from nanoCT datasets, illustrating age-associated changes in follicle number, localization, and health/atretic status. Bar, 250 µm.

### A stage-resolved mathematical model to uncover rules underlying symmetric follicle organization

Next, we investigated whether the maintenance of symmetric follicle spatial organization might be explained by underlying mathematical rules governing follicle growth and elimination. The model was applied first to the whole dataset of follicle counts, and then separately to the A-P and D-V halves, adopting a discrete finite-state linear framework to describe progression from secondary T4 (the first stage containing both healthy and atretic follicles) to preovulatory T8. Rates were assessed using follicle counts from 19 dpb to 4 mpb, a temporal window encompassing all types including those with atresia. Applying the linear equation described in Method Details, we developed a final system of equations (Figure 7A) that models follicle dynamics during transition between consecutive follicle types, graphically summarized in Figure 7B. To estimate the follicles’ transition rates, i.e., maturation to the next stage, retention within the current stage, or elimination, we employed Least Squares Estimation, validated with 10 repetitions of stratified k-fold cross validation and residual (error) analysis for each of the 10 sets of fitted parameters. The model identified stable transition rates between follicle stages from prepuberty to adulthood. The estimated transition rates revealed clear stage-specific patterns of follicle progression and elimination. Once recruited, the majority of T4 (61%) and then T5 (72%) mature to T6 (Figure 7C, D), resulting in an accumulation of this follicle type, which is consistently the most abundant (Figure 3C; Figure 5D; Figure S2A). This accumulation is also explained by the markedly lower recruitment of follicles progressing beyond T6, with only 17% reaching T7, and by the low elimination rate of atretic T6 (13%). Statistical analysis confirmed that the residuals in the test set followed a normal distribution (9 out of 10 repetitions) and that no significant bias was observed (10 out of 10) (Figure 7E). The consistent reduction in follicle number limited the applicability of the mathematical model beyond 4 mpb. To investigate whether the dynamics of follicle growth and elimination are preserved when the ovary is divided along its anatomical axes, we fed the model separately with follicle counts from the A-P or D-V halves. The estimated transition rates in A-P or D-V portions were statistically indistinguishable (Figure 7F), supporting a spatially symmetric distribution of follicles since its establishment in prepuberty. In summary, these results suggest the existence of a mutual relationship between the dynamics of follicle growth and elimination and the symmetric follicle distribution inside the ovary.

**Figure 7.**
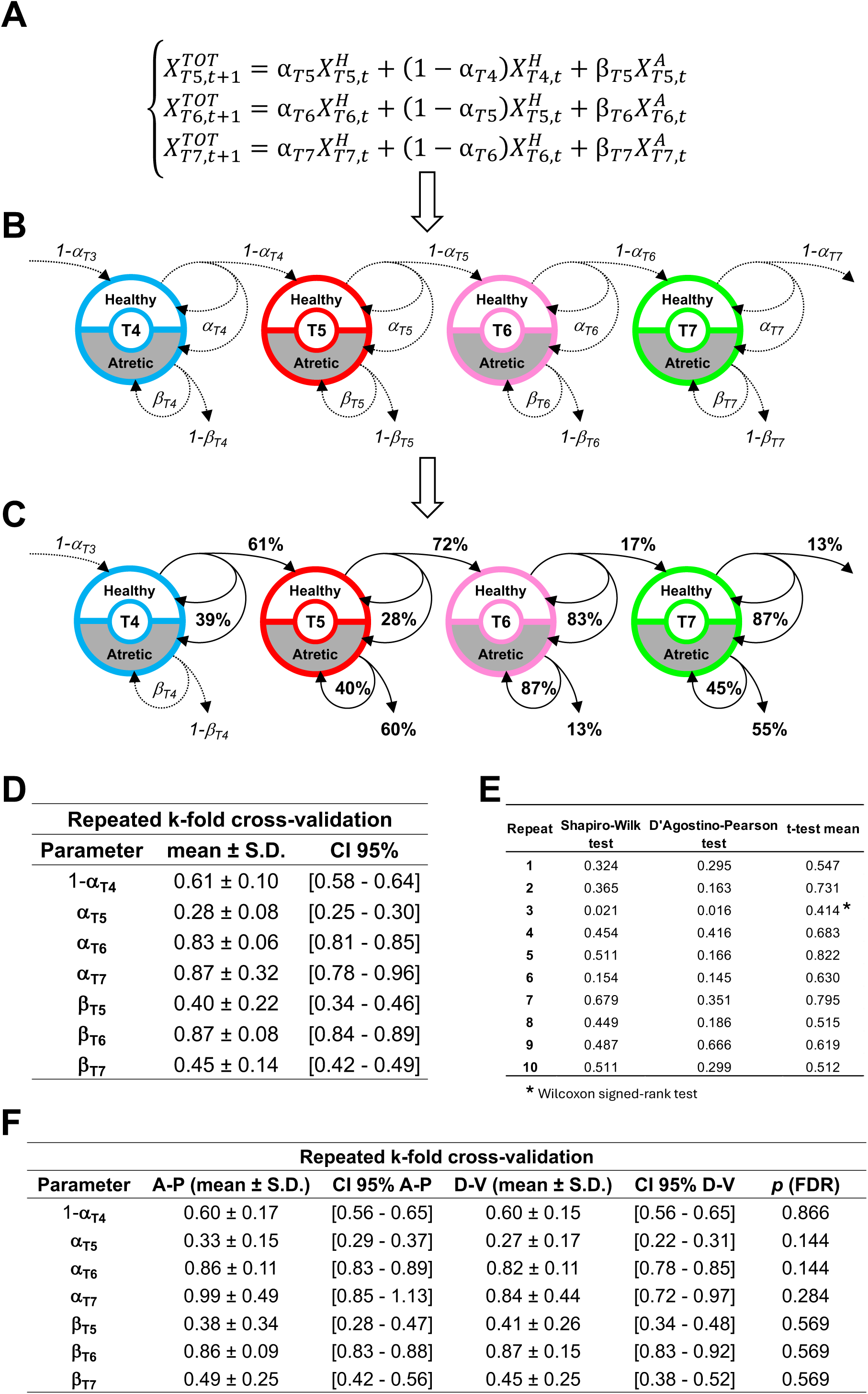
Mathematical modeling of follicle dynamics underlying symmetric organization (A) System of equations governing follicle state transitions. α_Ti_: healthy follicle retention rate; 1– α_Ti_: follicle growth rate; β_Ti_: atretic follicle retention rate. For details see Method Details. (B) Diagram of the finite-state linear model describing transitions of healthy and atretic follicles from T4 to T7. 1-β_Ti_: atretic follicle elimination rate. (C) Estimated transition rates derived from Least Square Estimation. The model highlights T6 as a key bottleneck, with high recruitment from T4 and T5 and limited progression or elimination. (D) Model validation through repeated five-fold cross-validation (n = 10). (E) Results of the residual analysis. (F) Five-fold cross validation of the model applied on anterior-posterior (A-P) or dorsal ventral (D-V) counts, respectively. *p* (FDR) is the result of the comparison between A-P *vs.* D-V estimated parameters.

## DISCUSSION

This study establishes a comprehensive 3D atlas of the mouse ovary that quantifies the spatial and temporal dynamics of primary to preovulatory follicles across infancy, prepuberty, adulthood, and aging, providing a basis to uncover organizing rules of ovarian function. We demonstrate that mouse ovarian follicles, regardless of developmental stage (T3 to T8), health status (healthy or atretic), or oocyte’s transcriptional activity (NSN or SN), are arranged in a quantitatively symmetric spatial pattern along the A-P and D-V axes. The symmetric follicular architecture and numerical order are established early in prepuberty (Video S7; Figure 3D) and maintained throughout life (Video S8; Figure 5E), underlining the stabilization of follicle dynamics despite the progressive decline in follicle number and the architectural remodeling imposed by CL and by the medullary accumulation of ZPRs.

Symmetry is already detectable at the primary T3 stage where, in the absence of vascularization [16], spatial distribution might be maintained by intrinsic organizational cues such as local biomechanical properties and molecular factors [17–20], supporting diffuse initial follicle growth [1]. Notably, during prepuberty and adulthood, the mean number of T3 remains in the range of 20-30, indicating that this represents the pool of primary follicles continuously feeding the folliculogenetic process at each cycle. Since the number of secondary T4 consistently matches or slightly exceeds that of T3, which are never atretic, this pattern suggests that most primary follicles progress to the next stage.

Secondary T4 are the first to be abundantly eliminated by atresia. At this stage theca cell layers begin to form, and stromal capillaries reorganize around the follicle periphery, establishing the first continuous vascular sheath that provides the metabolic and endocrine support essential for follicle survival, maturation or elimination [21–23]. As folliculogenesis proceeds, preantral T5 and particularly early antral T6 are the most prone to atresia, with the latter accounting for 30 to 50% of all atretic structures in the gonad, most of which accumulate toward the ovary’s center. This pronounced peak of atresia occurs during the transition from FSH-independent to FSH-dependent folliculogenesis [24,25], when follicles are rescued by FSH-driven activation [26].

These observations point to conserved rules of growth and elimination already established in early prepuberty, with 19 dpb marking a turning point when both healthy/atretic follicles are present. Consistently, our data indicate that no hallmarks of atresia (Figure 1D) [3] are detectable at 16 dpb, but the first five out of seven appear by 19 dpb, suggesting their onset within a three-day window. To refine this estimate, a more detailed day-by-day analysis of these morphological hallmarks, combined with immunodetection of molecular markers of atresia, would allow more precise timing of the events, from the earliest molecular degeneration signals to the execution of the atretic process. The sixth hallmark emerges at 22 dpb, and, in adults, atresia culminates with the presence of ZPRs. At 25 dpb, ∼50% of follicles are atretic, and about one-third continue to undergo atresia in adulthood and aging. By 9 months, the ovary begins its follicle numerical decline, although their symmetric distribution persists. Consistent with the depletion of the ovarian reserve [27–29], follicle numbers decrease by about half across all types and then stabilize, marking a significant age-related reduction of ovarian activity. Alongside this decline, we uncovered an age-dependent accumulation of ZPRs, extracellular glycoprotein matrices persisting after follicular atresia. They are absent in prepuberty but emerge by 2 mpb and reach a peak at 12 mpb with a ten-fold increase. Our data are consistent with ZPRs considered as long-lived structural markers of past follicular activity [28], shaping the aged ovary’s microenvironment and serving as quantifiable records of cumulative reproductive history. Their preferential retention in the medulla is a further indication of regionally biased degradation of atretic follicles.

Altogether, these findings highlight a balance among follicle input, growth, and elimination from late prepuberty to adulthood, suggesting a dynamic equilibrium. Leveraging quantification of follicle types across ovarian development, we formulated a stage-resolved mathematical model that formalizes folliculogenesis as a discrete finite-state system. The model uncovers two key organizational aspects operating from prepuberty to adulthood. It identifies a stage-specific bottleneck at T6, where high recruitment from T4-T5 combines with limited progression and persistence of atretic T6, resulting in their marked accumulation. Furthermore, it supports the existence of a symmetric ovarian architecture, in which follicles spatial distribution emerges from rules of growth and elimination conserved from prepuberty to adulthood. A key refinement of our mathematical model could include the subset of morphologically healthy follicles that are already committed to degeneration and would be detected as atretic in later counts. Moreover, a further integration of data on vascular remodeling, biomechanical changes and endocrine feedback could provide the model with functional multi-scale descriptive power under physiological and pathological conditions.

Within this symmetric architecture, oocyte’s nuclear organization emerges as both a marker and a stage-specific signature of follicle fate and developmental competence. Our nanoCT-confocal approach integrated structural and functional hallmarks of follicle status with oocyte’s transcriptional activity in the 25 dpb ovary, when the gonad has fully established its spatial and numerical follicle organization. The results demonstrate that follicles enclosing transcriptionally active NSN or inactive SN oocytes [13,14] are uniformly distributed across spatial axes, with nuclear states marking follicle fate in a stage-specific manner and linking chromatin organization to growth, atresia, and developmental competence. NSN chromatin characterizes healthy growing follicles at earlier stages but is absent in preovulatory healthy oocytes. In contrast, SN chromatin marks follicle degeneration in preantral and early antral follicles but is a feature of developmental competence in preovulatory T8, providing experimental support for a dual significance of this nuclear reorganization event previously hypothesized [13]. Taken together, these results underscore the importance of interpreting NSN/SN status within a stage-specific developmental framework. Importantly, by integrating this information, the 3D digital model provides a clear visualization of follicles in a state of atretic (NSN-Atretic and SN-Atretic), maturational (NSN-Healthy) or developmental (SN-Healthy) competence (Video S5).

Our non-disruptive nanoCT anatomical reconstruction provides a foundation for integrating a variety of functional information, exemplified here by the use of NSN/SN markers. Immunolabelling and spatial omics profiling [30] could provide insights into the molecular landscape of specific cells in their natural 3D environment, linking gene activity to ovarian architecture. Further integration with the biomechanical properties of the ovarian tissue [1] and the spatial-temporal reorganization of blood vessels would clarify how physical forces and vascular networks shape follicle survival, maturation, and degeneration. Together, these approaches would expand the 3D ovary atlas into a multi-scale framework for understanding ovarian physiology in health and disease.

In conclusion, the paper presents the 3D story of the developing ovary from birth to old age. Our study reveals the dual logic of ovarian organization: a robust spatial order coupled with selective stage constraints sustaining reproductive efficiency throughout the lifespan. These principles of spatial organization are not unique to the ovary but reflect a broader developmental logic found in other organs, in which our experimental pipeline could be applied, such as the lung (arborized bronchial tree; [31]), kidney (ureteric branching; [32]), intestine (crypt-villus units; [33]) and brain (branching arrangements; [34]), where reproducible geometric patterns are established early in development and integrated with a stromal-vascular framework.

Using nanoCT-based 3D reconstruction in different species will allow comparisons of ovarian structure, showing whether features like symmetry, stage bottlenecks, and continuous follicle input are shared or unique to each species’ reproductive strategy. These cross-species studies could improve our understanding of ovarian evolution and support applications in animal reproduction, fertility preservation, and conservation. The spatially symmetric organization of follicles, independent of atretic remodeling and aging, suggests the existence of an evolutionarily optimized developmental program in the mouse ovary. This organization supports coordinated folliculogenesis and balanced follicle recruitment across the ovary, sustaining the high ovulatory output characteristic of reproductive strategies that favor fast reproduction with large litters.

## METHODS

### Animals and Reagents

All experimental procedures adhered to the principles of European Union Directive 2010/63/UE and Italian Legislative Decree No. 26/2014. Female CD-1 mice (Charles River Laboratories, USA) were maintained under controlled conditions of 21 °C, 60% air humidity and a 12:12-hour light/dark cycle. All reagents were purchased from Sigma-Aldrich (Merck).

### Ovaries isolation and fixation

Three mice per age (1, 4, 7, 10, 13, 16, 19, 22, and 25 days post birth, and 2, 4, 9, 12, 14, 16, and 18 months post birth) were used to isolate 48 left and 48 right ovaries, with extra-ovarian annexes removed. Left ovaries, destined for nanoCT analysis, were fixed in 4% paraformaldehyde (PFA) overnight at 4°C with stirring, and the excess of fixative was removed by washing in distilled water (dH_2_O) for 24 hr. In parallel, right ovaries, destined for histological analysis, were fixed in Bouin’s solution (15 mL of picric acid, 5 mL of formalin and 1 mL of glacial acetic acid), and the excess was washed out with 70% ethanol.

### Contrast agent treatment and ovary embedding

Following PFA fixation, left ovaries were treated with 25% Lugol solution (2.5 gr potassium iodide and 1.25 gr Iodine in 100 mL of dH_2_O) at room temperature with stirring for a period varying depending on their size: 1 min for 1-4 dpb ovaries, 2 min for 7 dpb, 4 min for 10 dpb, 3 hr for ovaries from 13 dpb to 18 mpb. Excess contrast agent was removed by washing ovaries in dH_2_O for 15 hr (prepubertal) or 24 hr (adult and aging). After dehydration ovaries were embedded in paraffin blocks to be subjected to tomographic imaging.

### NanoCT images acquisition and pre-processing

Samples were scanned using the EasyTom XL Ultra 230 160 µ/nCT scanner (RX Solutions, Chavanod France) equipped with a Hamamatsu nano-focus transmission X-ray tube featuring a 1 μm-thick tungsten target on a diamond window and operated with a lanthanum hexaboride (LaB_6_) cathode. Projections were acquired using a CCD detector coupled to a Gadox (P43) scintillator screen (4008 x 2676 pixels) at 45 kV, 130-165 µA current and 1.85 sec exposure time, yielding a nominal voxel size of 250-850 nm. NanoCT datasets were reconstructed in TIFF format, maintaining the same voxel size, using X-ACT software (version 1.1, RX Solutions, Chavanod France). The reconstruction employed a filtered back-projection algorithm with ring removal and Hann apodization filters to suppress acquisition artifacts. Phase retrieval was carried out according to the single-material Paganin method with a 10% correction factor, equivalent to a-6 dB attenuation at 0.63 f_Nyquist_ (≈0.315 cycles px⁻¹), providing enhanced phase contrast while preserving spatial resolution. Image visualization and subsequent analysis were carried out using Fiji software [10].

### Follicle classification, counting and 3D localization

Follicles were classified as T3, T4, T5, T6, T7, T8, CL or ZPRs according to their morphological features. Follicles classification criteria were presence of the oocyte and ZP, presence and size of the antrum, and thickness and number of follicle cell layers surrounding the oocyte, which define the follicles diameter [2,9]. Follicles were classified as atretic if they exhibited additional parameters such as ooplasmic cavities or complete oocyte degeneration, granulosa cells detaching from the ZP, or the presence of apoptotic or fibroblast-like cells in the antral cavity [3]. ZPRs are characterized by a small, completely degenerated follicle containing only a collapsed, empty ZP.

Using Fiji’s *ROI Manager* tool, each follicle was annotated with a colored dot according to the follicular type (T3, purple; T4, light blue; T5, red; T6, pink; T7, green; T8, orange; CL, yellow; ZPRs, bordeaux) and manually tracked across consecutive nanoCT sections to ensure accurate identification and counting. Atretic follicles were further distinguished by adding a black central mark within the color dot. To localize and count follicles, CLs, and ZPRs, we used a spatial reference system based on the organ’s anatomical orientation, rotating each nanoCT image stack to achieve the final alignment shown in Figure S6. When counted, follicles were assigned to either the dorsal, ventral, anterior, or posterior region, depending on whether more than 50% of their volume belonged to one or the other.

### Histological slides preparation and analysis

After embedding in paraffin, right Bouin’s fixed ovaries were sliced into 10 µm thick serial sections with a RM2125RT rotating microtome (Leica Biosystems). Sections were stained with Mayer’s hematoxylin and Y eosin or with PAS-hematoxylin and analyzed using a DM6B microscope (Leica Microsystems) equipped with a LAS X Navigator stitching software (Leica Microsystems).

### nanoCT-confocal imaging co-registration

Three 25 dpb ovaries previously analyzed with nanoCT were sectioned at 20 µm thickness and mounted on glass slides coated with poly-L-lysine. Iodine residuals from Lugol staining were removed by treating sections with 5% sodium thiosulfate for 5 min at RT. Then, sections were incubated overnight at 4°C with 0.5 µg/ml 4′,6-diamidino-2-phenylindole (DAPI) in 1X PBS and mounted in an antifading medium (1:1 mixture of DABCO and 0.5 µg/ml DAPI). Confocal images were acquired using a TCS SP8 microscope (Leica Microsystems) equipped with a 40x objective. High-resolution stacks with 1 µm z-step of the entire section thickness were obtained using the *Navigator* stitching function of the LAS X software.

LIF images from individual stacks were concatenated into a single dataset using the *Concatenate* tool in Fiji. The resulting confocal dataset was used as reference to manually co-register nanoCT images of the same ovary using the *Interactive Stack Rotation* plug-in in Fiji (Figure S4).

Each oocyte was classified according to chromatin organization and annotated through the *ROI Manager* tool as follows: green tag, Surrounded Nucleolus (SN), characterized by a continuous ring of heterochromatin encircling the nucleolus; red tag, Not Surrounded Nucleolus (NSN), defined by heterochromatin dispersed as discrete spots throughout the nucleoplasm. Additionally, oocytes presenting a degenerated or condensed chromatin were considered unclassifiable and tagged in white. Following annotation, the nanoCT dataset was realigned to the original D-V axis, allowing the integration of information on oocyte chromatin organization, associated with its transcriptional activity, with previous data on follicle type. Red, green, or white squares were added using the *ROI Manager* tool in Fiji drawing their contours around the follicle type tag.

### Follicle tag 3D coordinates extraction

Follicle spatial localization was determined with reference to the centers of both the ovary and the follicles. A custom Python script automatically extracted the x, y, z coordinates of each colored tag center from annotated nanoCT image datasets. The resulting data were then sorted into distinct.csv files based on follicle type, atresia, or chromatin organization, and subsequently imported into Blender software.

### Digital models of the ovary using Blender

Blender version 4.2 was used to model the ovaries. To work with micrometer scale units, in ‘Scene Properties’ set the ‘Unit System’ with ‘Metric’ and ‘Unit Scale’ at 0.000001. In the ‘View Tab’ set the ‘Focal Length’ at 50 mm, the ‘Clip Start’ at 1 µm’ and the ‘End’ at 50,000 µm. To import the csv files, Blender needs a non-native spreadsheet import file (https://extensions.blender.org/add-ons/spreadsheet-import/) which needs to be added to the ‘Blender Preferences’ in ‘Themes’. Csv files containing x, y, z coordinates, follicles size and color are imported with the ‘Scripting’ function thanks to a first custom-made Python script. A second Python script automatically sectioned the ovary and categorized follicles along the A-P and D-V axes. As with expert-based annotation, the script assigned follicles to hemispheres based on whether more than 50% of their volume lay on one side of a given plan and exported morphometric data as csv files for downstream quantitative analysis. A third Python script generated an ellipsoid reference structure centered to the ovary’s center and representing the inner 25% of the gonad’s volume and colored follicles grey when located within this region.

### Mathematical model

Folliculogenesis was represented as a discrete finite-state linear model, where each follicle occupied a specific maturational state at a given time point. The temporal evolution of the system was modelled in discrete steps, from 19 dpb to 4 mpb. This interval corresponds to a developmental window in which all follicular states from T4 to T7, as well as atretic follicles, are observed. The temporal steps were not uniformly spaced, thus an alignment across ages was performed using Dynamic Time Warping.

PyCharm was the integrated development environment used to write the custom Python codes used. At each time step t (i.e., 19, 22 and 25 dpb and 2 and 4 mpb) and for each follicle type (i.e., T5, T6 and T7), the number of follicles in state 𝑇_i_ was defined as the sum of healthy follicles (𝑥*^H^_T,i,t_*) and atretic follicles (𝑥*^A^_T,i,t_*), such that X*_T,i,t_*=𝑥*^H^_T,i,t_*+𝑥*^A^_T,i,t_*, for a total of 15 data points.

For a more robust parameter estimation and model validation, the sample size for was increased exploiting the A-P and D-V spatial symmetries of the ovary, resulting in a dataset of 60 data points, equally distributed across the follicular states T5, T6, and T7.

The number of follicles in the subsequent temporal step 𝑥*_T,i,t+1_* was modelled as a linear equation:

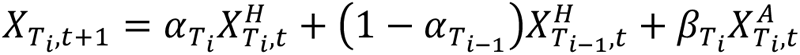

where 𝛼*_Ti_* represents the rate of follicles remaining in state 𝑇_i_ (either healthy or degenerated to atretic); 1-𝛼*_Ti-1_* represents the rate of 𝑇*_i-1_* follicles maturing into state 𝑇_i_; and 𝛽*_Ti_* represents the rate of atretic follicles not yet eliminated and therefore still observed.

To assess the 𝛼*_Ti_* and 𝛽*_Ti_* parameters that best minimize the difference between observed and modelled follicle counts, a Least Squares Estimation method was used. The model was validated by performing repeated stratified k-fold cross-validation (k = 5, repeated for n = 10 times with different random seeds to assess variability and robustness), ensuring that each fold preserved the class distribution (i.e., equal representation of the three follicle states). Residuals were computed for each of the 10 parameter sets obtained from the cross-validation procedure.

## Statistical analyses

Statistical analyses were performed using the IBM SPSS Statistic software. Normal data distribution was assessed with the Shapiro-Wilk test. For normally distributed data, one-way ANOVA was performed, followed by pairwise t-tests. For non-normally distributed data, the Kruskal-Wallis test was applied, followed by pairwise Mann-Whitney U tests with Bonferroni correction for multiple comparisons. For the mathematical model, the normal distribution of residuals (error) in the test sets of each repeat of the cross validation, was assessed applying both the Shapiro-Wilk and the D’Agostino-Pearson tests. Depending on distributional properties, either one-sample t-test (for normally distributed data) or Wilcoxon signed-rank test (for non-normal distributions) were employed to verify the absence of systematic bias in the model. Normal distribution of estimated parameters in A-P and D-V regions was assessed using the Shapiro-Wilk test. Comparisons were performed using tests for independent samples: independent t-test for normally distributed data and Mann-Whitney U test for non-normally distributed data. p-values were adjusted for multiple comparisons using the Benjamini-Hochberg false discovery rate (FDR) method.

---

**Figure S1.**
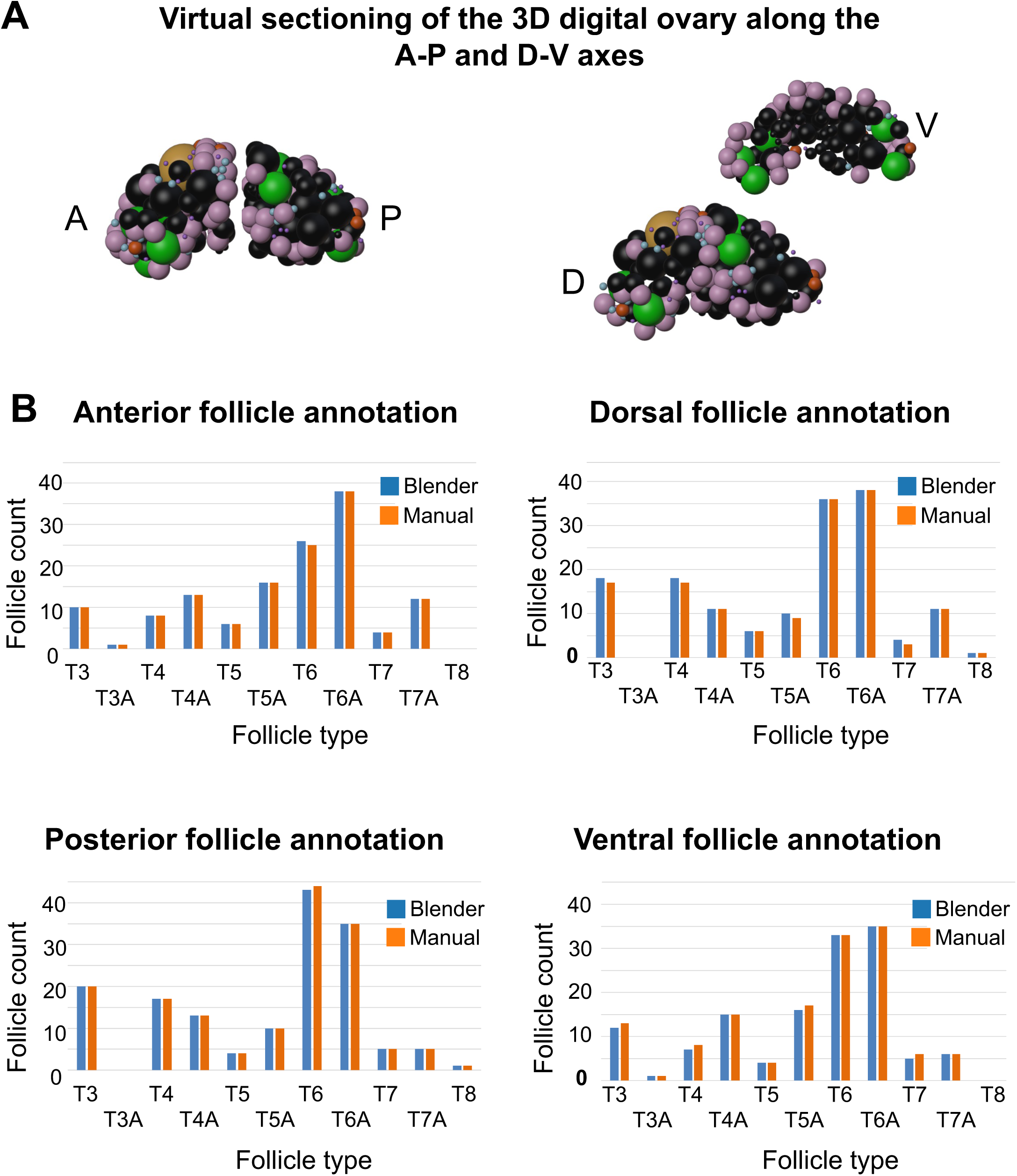
Validation of the 3D digital ovary model by comparison with expert-based annotation (A) Automated follicle count and localization using a Blender-based pipeline was validated against expert annotation by comparing counts along the A-P and D-V axes. As with expert-based annotation, the script assigned follicles to hemispheres based on whether more than 50% of their volume lay on one side of a given plan and exported morphometric data as csv files for downstream quantitative analysis. (B) Side-by-side comparison of follicle number and distribution in a single ovary obtained from automated (Blender-based) or expert-based annotation showing high concordance.

**Figure S2.**
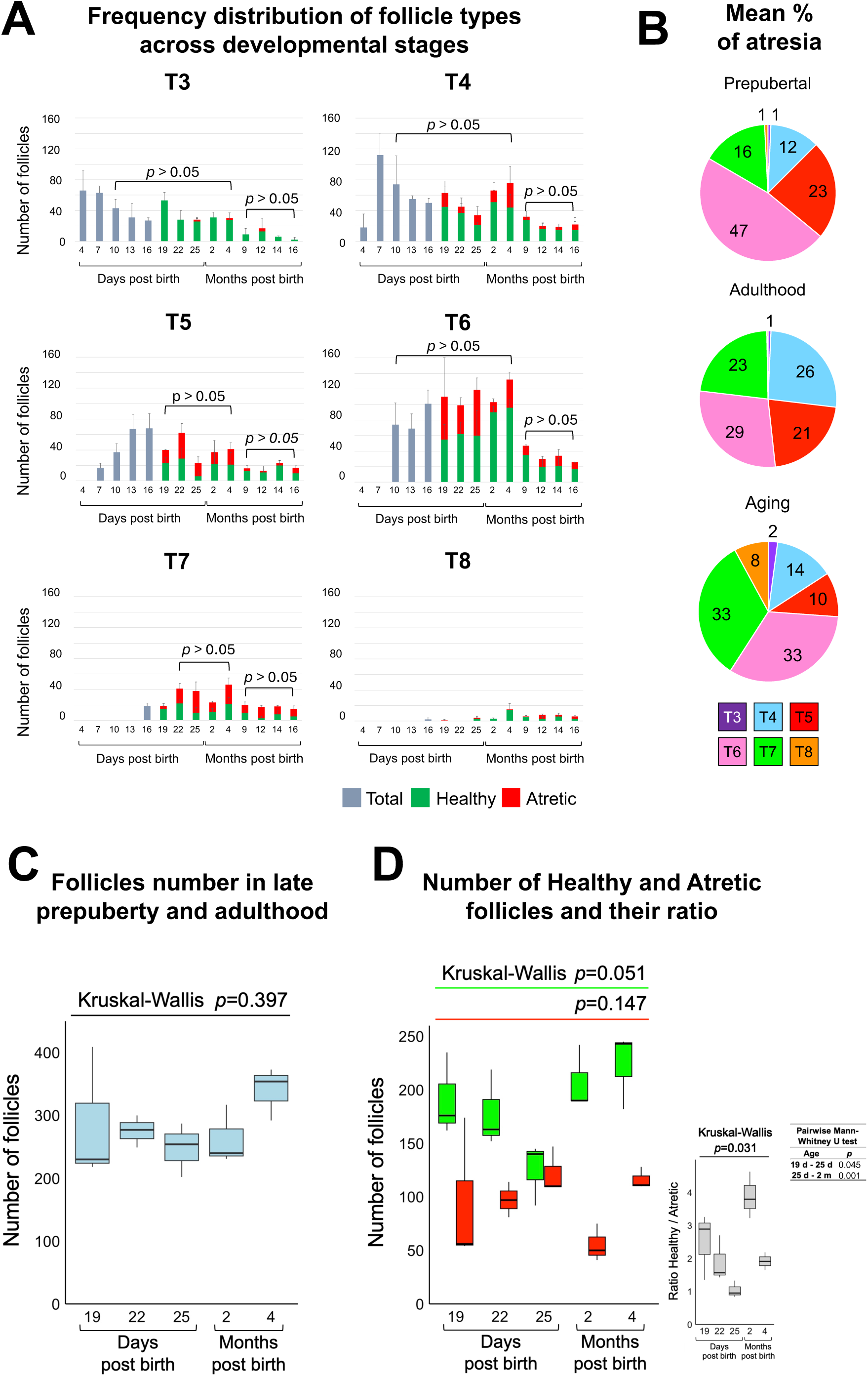
Changes in follicle number and viability across lifespan (A) The number of T3, T4 and T6 follicles remains constant from 10 dpb to 4 mpb; that of T5 follicles from 19 dpb to 4 mpb; and that of T7 follicles from 22 dpb to 4 mpb. Following a decline in number, from 9 to 16 mpb T3-T7 follicles remain quantitatively stable. (B) Frequency of T3-T8 atretic follicles in the prepubertal, adult and aging ovary. (C) Box plot of the total follicle number during late prepuberty (19-25 dpb) and adulthood (2-4 mpb), showing a stable number of follicles. (D) Box plots of the number of healthy and atretic follicles and their ratio, showing a fair number of healthy and atretic follicles, but a significant increase of the healthy population comparing the prepubertal 25 dpb with the adult 2 mpb ovary.

**Figure S3.**
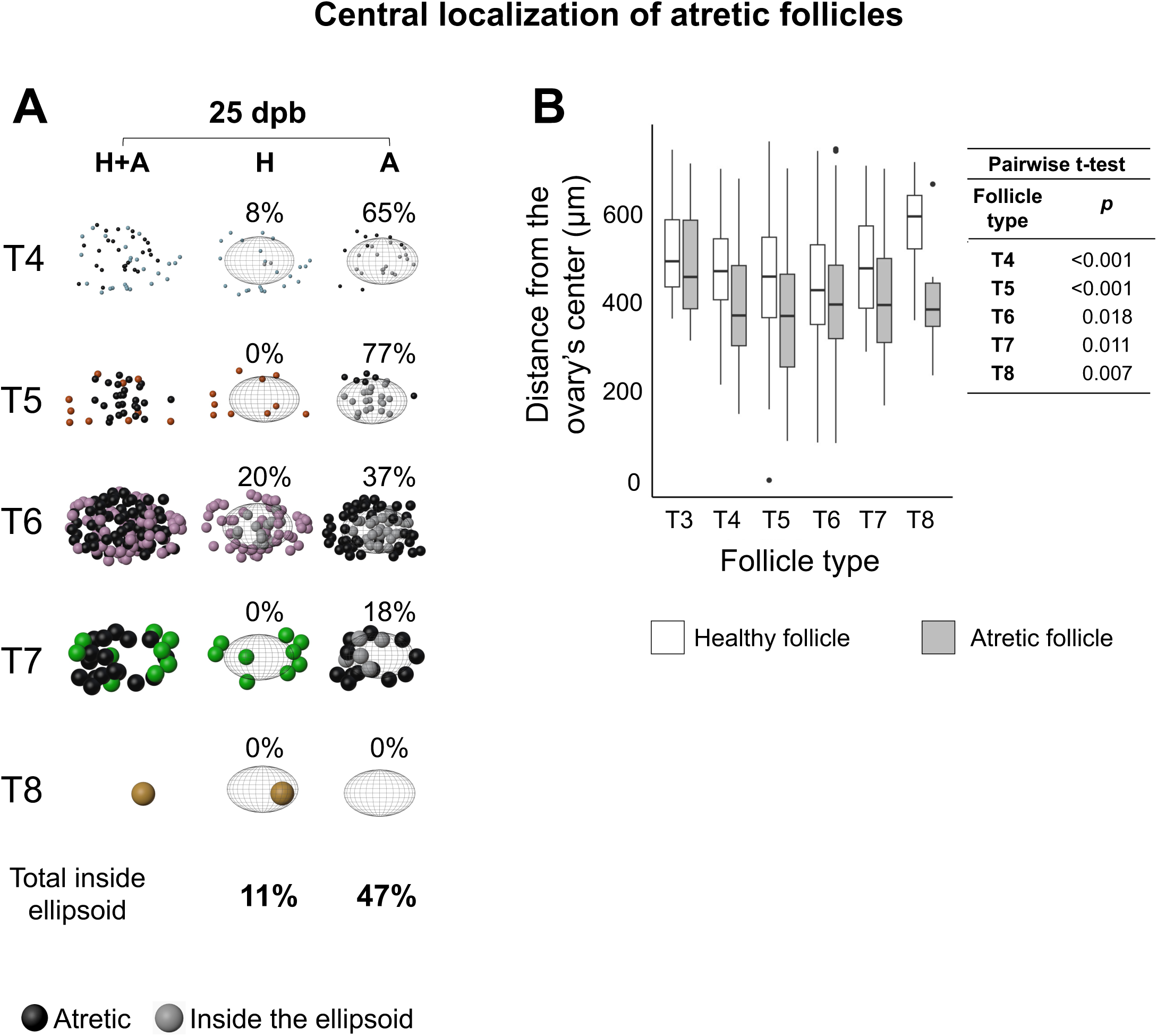
Atretic follicles are preferentially localized toward the ovarian center during late prepubertal development (A) Spatial analysis of follicle localization in a representative 25 dpb ovary using a centered ellipsoid reference model covering the inner 25% volume, selected because smaller volumes captured too few follicles, as the medulla (∼13.5% in adults; [35]) progressively fibroses and becomes follicle-poor with age. Atretic follicles (A) were more frequently located within the central ellipsoid region (in grey) than healthy follicles (H). (B) Quantification of the distance from the ovary center for healthy and atretic follicles across T3-T8 stages. T4-T8 atretic follicles were significantly closer to the center than their healthy counterparts.

**Figure S4.**
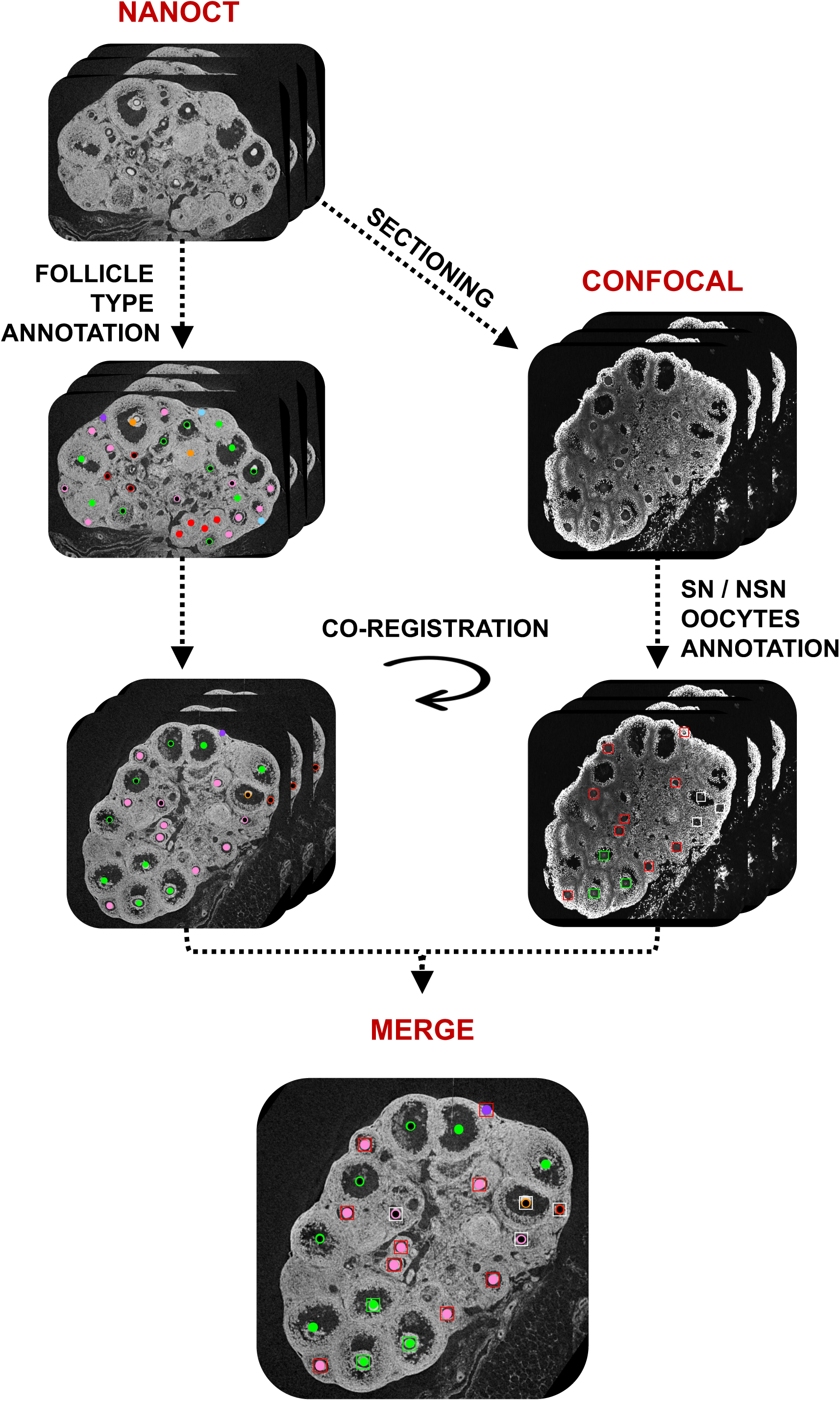
NanoCT-confocal imaging pipeline Following nanoCT imaging of 25 dpb ovaries, follicle types are annotated in the resulting datasets. In parallel, the same ovaries are sectioned into 20 µm-thick slices, stained with DAPI and analyzed by confocal microscopy to annotate follicles based on SN or NSN oocyte’s chromatin organization. Then, image co-registration enables the merge of both type of information.

**Figure S5.**
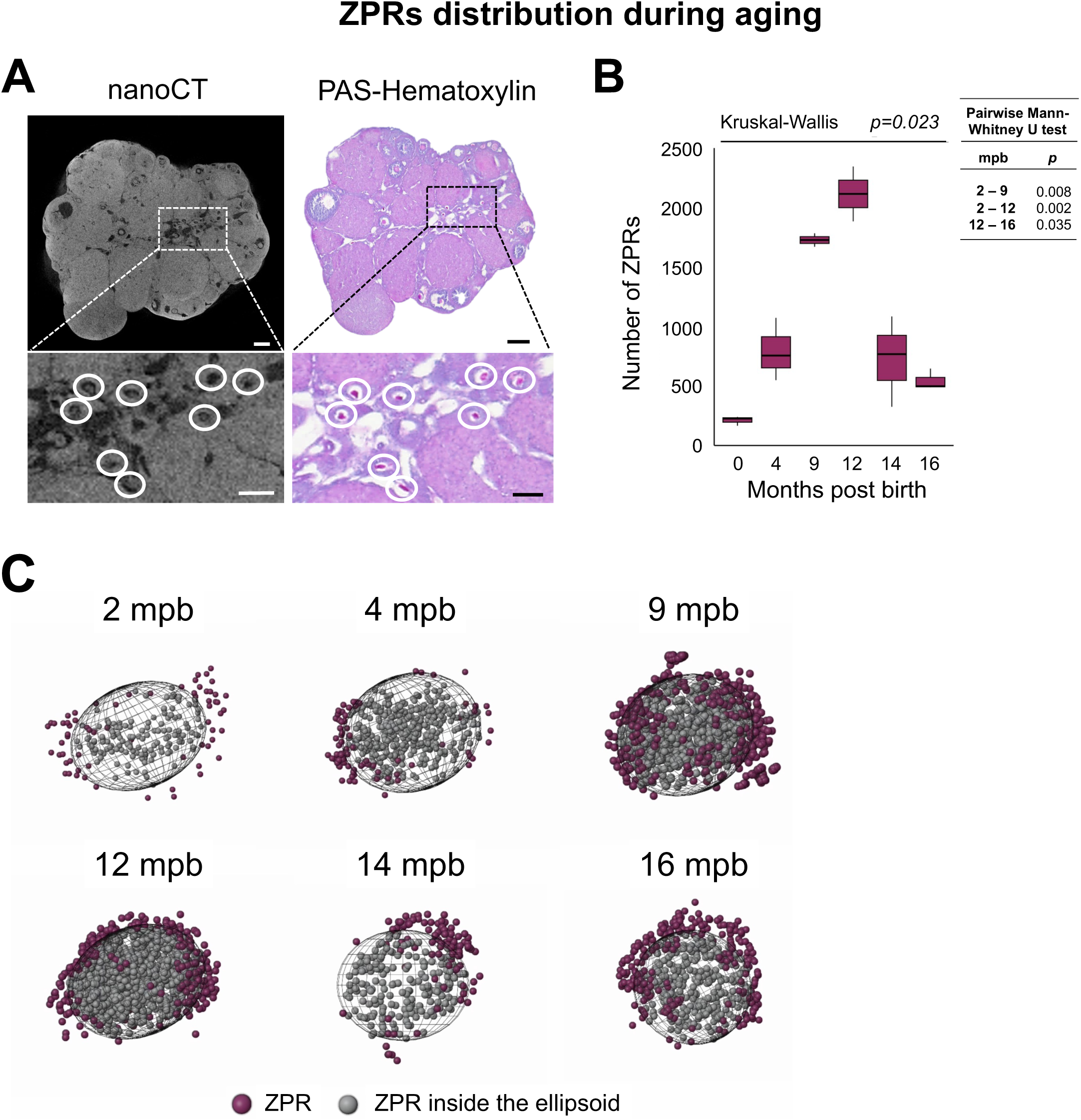
Age-associated spatial dynamics of zona pellucida remnants (A) Co-registration of nanoCT and PAS-hematoxylin histology [28] show the colocalization of glycoprotein-rich ZPRs. ZPRs were absent in prepubertal stages and first detected at 2 mpb. (B) Quantification of ZPRs across aging ovaries reveals a statistically significant ∼10-fold increase by 12 mpb followed by a decline at later ages. (C) Spatial analysis of ZPRs localization in representative 2-16 mpb ovaries showing the progressive accumulation inside the reference ellipsoid, reaching a peak at 12 mpb.

**Figure S6.**
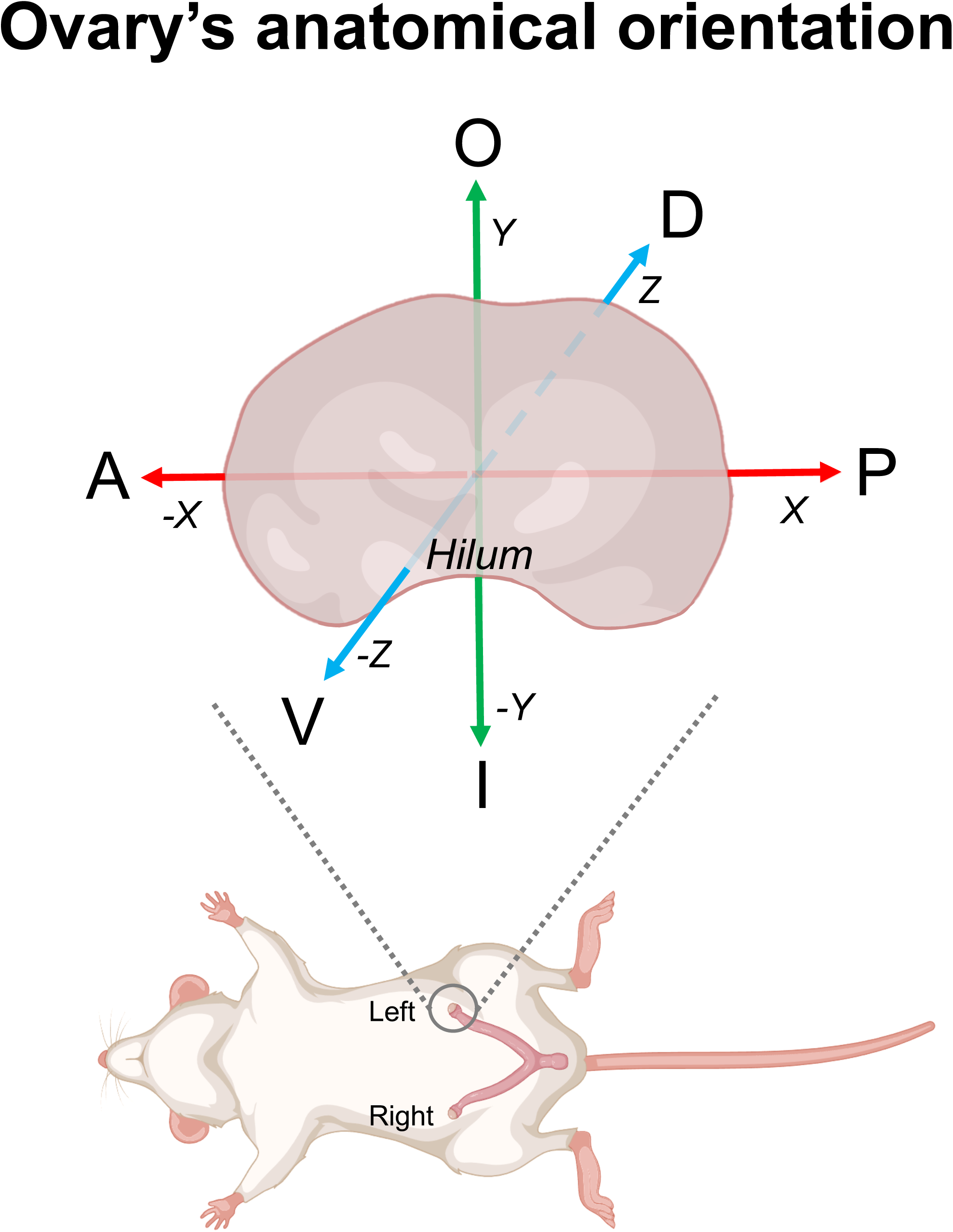
Anatomical orientation of the mouse ovaries used for nanoCT imaging.

